# Disregarding fish ageing method can compromise growth parameter comparability across studies

**DOI:** 10.1101/2025.05.13.653728

**Authors:** Elyza Pilipaityte, Eglė Jakubavičiūtė, Žilvinas Pūtys, Asta Audzijonyte

## Abstract

Reliable and comparable growth estimates are critical for accurate comparative studies, robust stock assessments, and effective fisheries management. Many studies compare fish growth Von Bertalanffy parameters (growth coefficient *K* and asymptotic length *L*_∞_) across species, populations, or time periods. These parameters are often obtained from literature or FishBase without accounting for ageing method used to estimate size-at-age, despite known differences in age estimates from different structures. In this study, we use four species as a case study and once again demonstrate that fish ageing based on otoliths and scales can produce significantly different estimates of *L*_∞_ and *K*. We then analyse available *K* and *L*_∞_ records from FishBase and show that *L*_∞_ values derived from scales were, on average, 15% higher than those obtained from otoliths, while *K* values were 20% smaller (p < 0.01 in both cases). We strongly recommend that ageing methods be clearly reported in all growth studies and included in databases, that comparative analyses explicitly account for ageing method in statistical model structure, and that researchers consider methodological differences when interpreting growth parameter variations across studies.

## Introduction

Fish life history parameters, such as Von Bertalanffy (Von Bertalanffy 1938) growth curve parameters, are essential inputs for fisheries assessments or ecosystem models and are widely used in large scale comparative studies (Thorson, 2020; Helminen & Sarvala, 2021; Spence et al., 2021; Palomares et al., 2022). FishBase (Froese & Pauly, 2024) is the main depository of fish life-history parameters, and many studies extract them for multiple species and populations to compare growth rates or body sizes across geographic and temporal scales (van Denderen et al., 2020; Palomares et al., 2022). Von Bertalanffy growth parameters are obtained by fitting a three-parameter curve to length and age data, where ages can be determined using different ageing methods. Traditionally fish ages have been determined using scales, but more recently otoliths have been increasingly popular. Comparing Von Bertalanffy growth parameters estimated from different ageing methods could introduce biases, as age determinations from different structures (scales, otoliths, vertebra, opercula) are known to produce different results (Robillard & Marsden, 1996; Bostanci, 2008; Muir et al., 2008). Specifically, ages determined from scales often seem to overestimate fish age, which would lead to underestimation of growth and overestimated asymptotic lengths (Bertignac & De Pontual, 2007; Rittweg et al., 2023). Moreover, discrepancies between age estimates derived from otoliths and scales can vary with the age of the fish—scales may overestimate the age of younger fish while underestimating that of older individuals (Goeman et al., 1984; Robillard & Marsden, 1996; Rittweg et al., 2023). These biases highlight the importance of accounting for ageing methods in large-scale analyses, as done, for example, in Dortel et al. (2013). Nevertheless, the ageing method is often overlooked in interpopulation or interspecies comparisons. In fact, many growth parameter estimates reported in FishBase do not even specify the ageing method used, implicitly suggesting that methodological differences are considered minor relative to other sources of variation in fish growth and body size. This study was prompted by growth analyses of four common freshwater fish species in the brackish water Curonian Lagoon (Baltic Sea) and observation that scale and otolith-based ages were considerably different in some species. We then conducted a broader study of Von Bertalanffy growth parameters data available on FishBase and show that not accounting for the ageing method used could introduce systematic biases in comparative analyses.

## Materials and Methods

### Estimation of growth parameters from different ageing methods

This study was prompted by a broader analysis assessing long-term growth changes of common fish species in the Curonian Lagoon – a shallow and highly productive water body located in the central eastern part of the Baltic Sea (Pilipaityte et al., submitted after revisions). The broader study conducted both scale and otolith-based ageing on same fish individuals, to ensure comparability with the long-term data series, where both methods were used. In this study, we focus on four common fish species: bream (*Abramis brama* L. 1758), roach (*Rutilus rutilus* L. 1758), perch (*Perca fluviatilis* L. 1758), and pikeperch (*Sander lucioperca* L. 1758). A total of 182, 195, 206 and 102 individuals of each species, respectively, were aged using samples collected in 2023 (using standard gillnetting survey methods, see Pilipaityte et al., submitted after revisions).

To determine fish age, both scales and otoliths were examined. Scales were collected from above the lateral line on the left side of each fish, just behind the dorsal fin, and viewed under a binocular microscope. Otolith-based age estimates were obtained from transverse sections of sagittal otoliths, following Proctor et al. (2021). One otolith per fish was embedded in resin, cured for 24 hours, and sectioned using a diamond saw to produce ∼500 µm thick slices containing the core. Sections were mounted on microscope slides using resin and hardener. Annuli were counted under a binocular microscope (10–20× magnification) using both reflected and transmitted light to enhance visibility. To assess reading accuracy all otoliths were aged independently by two readers, and a random subset of scale data was also assessed by an independent reader.

Age from the two ageing methods (scales and otoliths) was then plotted and compared visually and for both methods we also estimated Von Bertalanffy growth curve parameters *L*_∞_, which is the asymptotic length (cm), *K* which is the growth coefficient, and *t*_0_ which is the theoretical age at which fish length is zero. These three parameters give the full growth function:

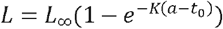

where L is the length-at-age. Parameters were estimated by fitting the Von Bertalanffy growth function to the age-length for scale and otolith data separately using the nonlinear least squares estimation, as implemented in the ‘vbFuns’ function in the R package FSA (Ogle, 2022).

### Comparative analysis of growth parameters from FishBase

Given that in our analyses *L*_∞_ and *K* estimates obtained from scales were consistently different than those obtained from otoliths, we then assessed whether a similar pattern could be found in other fish species. To do this, we extracted all available fish growth and ageing method parameters from FishBase (http://www.fishbase.org, last accessed July 2024) (Froese & Pauly, 2024) using the ‘rfishbase’ package in R (versdion 4.1.2; Boettiger et al. 2012). We also extracted additional information, such as the year of the study, location and other notes. The total amount of initial data included 13374 estimates of *L*_∞_ parameters for 2689 species. These data were screened to exclude: i) all *L*_∞_ estimates that were identified as doubtful by FishBase (1989 entries), ii) *L*_∞_ estimates that did not include the ageing method (3286 entries), iii) *L*_∞_ estimates obtained from structures other than scales or otoliths (4184 entries). For the remaining 3885 observations from 1283 species, most species had only a single *L*_∞_ estimate from one or the other ageing method, which did not allow for a statistical comparison. We therefore filtered the data further to include only those species that had at least four *L*_∞_ and *K* estimates from both scales and otoliths (at least eight estimates in total). This filtering gave us 220 fish growth parameters estimates for nine species (Fig. 1).

**Fig. 1:**
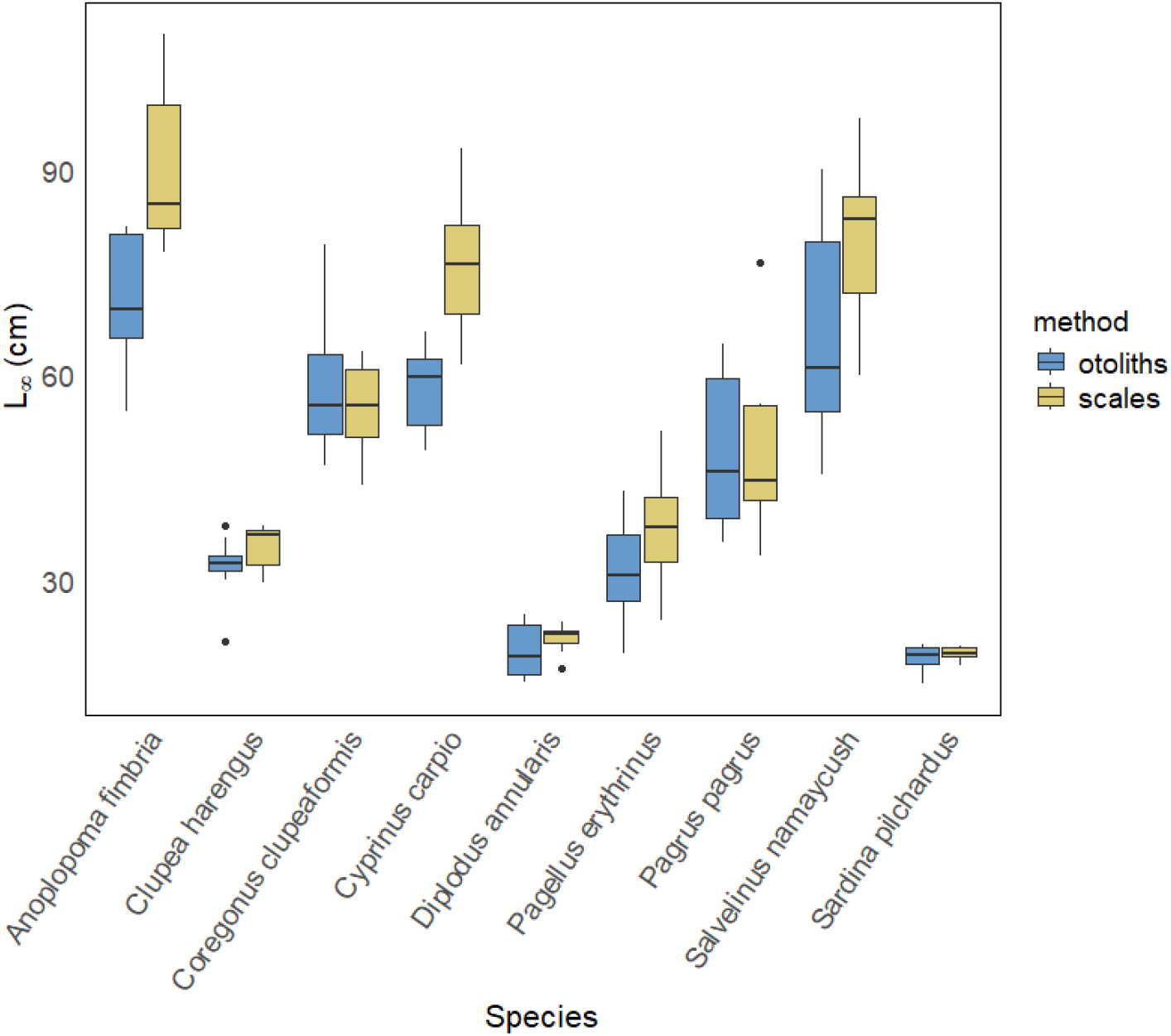
Species-specific comparisons of asymptotic length (*L*_∞_) estimated using scales versus otoliths from the dataset available on FishBase (where at least four estimates per method per species were available)

To assess whether *L*_∞_ and *K* estimates differed between the two ageing methods we used a linear mixed effect regression models (function *lmer*, library ‘*lme4’*; Bates *et al*., 2015). Specifically, *L*_∞_ and *K* was modelled as a function of the ageing method, while species were treated as a random effect to account for different absolute *L*_∞_ and *K* values across species. Our model was described as:

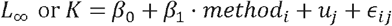

where *β*_0_ is the intercept, *β*_1_ is the coefficient for the fixed effect, *u*_j_ is the random effect for species *j, ϵ*_ij_ is the residual error for observation in species, assumed to be normally distributed with mean 0 and variance *σ*^*2*^. The model was fitted using maximum likelihood estimation. It was not possible to statistically test the effect of reporting year or location on growth parameter estimates, because there were too few observations per location and years. However, data exploration showed that there were no obvious differences in the years between the two ageing methods, as both scales and otoliths were used throughout the entire study period (see data, code and additional images on GitHub https://github.com/ElyzaPilipaityte/Linf_AgeingMethod).

## Results

### Large differences in growth parameter estimates from two ageing methods

In the first part of our analyses, comparison of otolith- and scale-based ages in four Curonian Lagoon fish species showed that ages and estimated Von Bertalanffy parameters from the two methods were substantially different, especially for perch and pikeperch (Fig. 2). The difference in age estimates between the two independent readers was generally within 0–1 year for both otoliths and scales. However, for bream, discrepancies were larger – ranging from 0 to 4 years in otoliths and 1 to 2 years in scales.

**Fig. 2:**
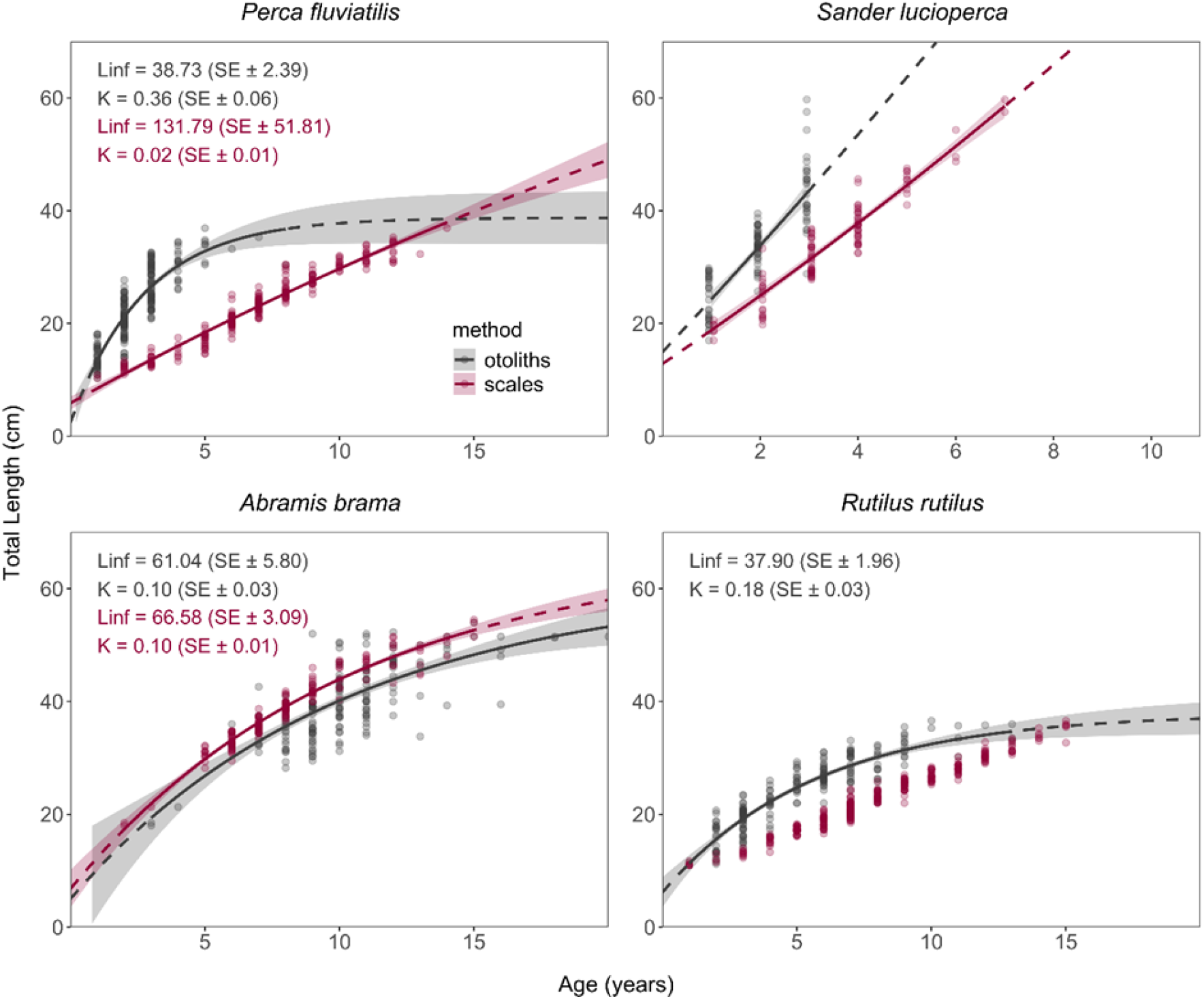
Von Bertalanffy growth curves (with 95% confidence intervals, shaded area) estimated for four fish species estimated from two ageing methods – scales and otoliths. The solid line indicates growth curves fitted to observed age and length data, while the dashed area indicates the extrapolation outside the observed range

In contrast, the differences between ageing structures used were much greater than those between readers. For perch aged at 2 years with otoliths, scale ages ranged from 3 to 8; for 3-year-old perch, ages determined by scales varied between 7 to 11. The difference between ageing structures for pikeperch was also obvious, with the oldest fish estimated to be 3 years old based on otoliths and 7 years old using scales. Age differences from scales and otoliths were smaller in roach, but scales were still giving 1–6-year older ages for some individuals compared to otoliths. For bream differences in determined ages were smaller although for some individuals there still were up to 4-year differences in ages determined from scales and otoliths.

For perch age data from scales led to unrealistic *L*_∞_ estimates (Fig. 2). Fitting growth curves and calculating *L*_∞_ for pikeperch was challenging for both ageing methods, because the population is overfished and age structure is strongly truncated (Jakubavičiūtė et al., 2024). Bream was the only species with no substantial differences in growth curves determined by the two methods, with *L*_∞_ estimates being similar between scales and otoliths.

### Analysis of available growth parameters from FishBase

Analysis of FishBase data revealed that ageing method alone resulted in a significant difference in *L*_∞_ and *K* estimates across studied species (p < 0.001 and p = 0.011 respectively). Specifically, *L*_∞_ values derived from scales were, on average, approximately 15% higher than those obtained from otoliths (Table 1), whereas *K* values were approximately 20% lower. Looking at individual species estimates, the trend was generally consistent, though the magnitude varied among species (Fig. 1), with scale-based *L*_∞_ estimates in e.g. *Anoplopoma fimbria, Cyprinus carpio*, and *Salvelinus namaycush* substantially exceeding those derived from otoliths. In contrast, species such as *Clupea harengus, Sardina pilchardus*, and *Diplodus annularis* showed smaller differences between methods.

**Table 1:** Linear mixed effect model coefficients indicating the effect of the ageing method (otoliths or scales) on the asymptotic length *(L*_∞_) and growth coefficient (*K*) estimates of nine fish species from FishBase. Marginal R^2^ indicates the proportion of total variance explained by the fixed effect, while conditional R^2^ shows the proposal of total variance explained by fixed and random effects. Most of the variation was explained by species identity due to different sizes of fish species

| *Coefficient* | *Estimate* | *95% CI* | *P-value* |
| --- | --- | --- | --- |
| $L_{\infty}$ | | | |
| Intercept | 45.66 | 31.31– 60.01 | <0.001 |
| Method [scales] | 6.92 | 4.10 – 9.74 | <0.001 |
| Marginal $R^2$ / Conditional $R^2$ | 0.021 / 0.846 | | |
| $K$ | | | |
| Intercept | 0.33 | 0.27 – 0.39 | <0.001 |
| Method [scales] | -0.07 | -0.13 – -0.02 | 0.011 |
| Marginal $R^2$ / Conditional $R^2$ | 0.031 / 0.138 | | |
| Observations | 220 |  |  |

## Discussion

Discrepancies between scale and otolith derived age estimates shown in this study are not new and have been reported before. Our findings are consistent with earlier studies, demonstrating that age derived from scales tends to overestimate *L*_∞_ and underestimate *K* (Lozano et al., 2013; Tyszko & Pritt, 2017; Rittweg et al., 2023), because ages in scales often appear to be older than they are in otoliths, especially in younger ages. However, opposite results have also been reported in some seabream species (Abecasis et al., 2008) and it is not necessarily clear whether a consistent bias should be expected. Ageing material has been identified as a key source of systematic bias in the estimation of growth parameters such as *L*_∞_ and *K* in dolphinfish (*Coryphaena hippurus*) (Chang & Maunder, 2012). Bertignac and De Pontual (2007) showed that age estimation errors can significantly influence estimates of fishing mortality and stock biomass in European hake (*Merluccius merluccius*). Similarly, inaccurate age estimates derived from scales have also led to underestimations of fishing mortality in Largemouth Bass (*Micropterus salmoides*) (Tyszko & Pritt, 2017), and to overestimations of maximum sustainable yield in northern pike (*Esox lucius*) from the southern Baltic Sea (Rittweg et al., 2023). In our study, the use of scale-based age estimates would likely have led to underestimation of fishing mortality for Curonian Lagoon perch, pikeperch, and roach. This is because scale-based data suggested that age composition of bream, roach and pikeperch includes a lot more older fish than otolith-based estimates. Although both scales and otoliths are commonly used ageing structures, otolith-derived age estimates are generally considered more reliable in many species (Goeman et al., 1984; Ashford et al., 2001; Schill et al., 2010). Given the overexploited status of pikeperch in Curonian Lagoon, the otolith-based estimates align more closely with the expected demographic structure and thus are likely to be more accurate.

While the difference between scale and otolith derived age estimates is not new, our study shows that not accounting for ageing source in comparative analyses of existing datasets can lead to substantial (15-20%) differences in parameter estimates. This will vary across species, where e.g. both methods produced similar outcomes in bream, but very different results in perch and pikeperch. Crucially, if different ageing methods are used, the resulting parameters may not be directly comparable, potentially undermining the validity of cross-study comparisons through space and time. This has significant implications not only for comparing growth estimates across studies or regions but also for stock assessments, where consistent and accurate age data are essential for reliable population modelling and management decisions. The average difference of 15-20% in *K* and *L*_∞_ parameters is large enough to lead to substantially different outcomes of, for example length-based spawner potential ratio (LB-SPR) stock assessment methods (Hordyk et al., 2015). Of course, cross-studies comparisons based on existing data on FishBase and other literature sources may suffer from other issues, such as inaccurate overall age estimates, as was also the case in our analyses for scale data and likely also for some otolith-based readings. Addressing such inaccuracies requires careful consideration already at the point of data collection and is not the goal of our research. The main point we make here is that despite known differences in age estimation from scales and otoliths, there is still insufficient emphasis on accounting for ageing structure effects when using literature, historical or FishBase data. Given that older studies more commonly used scales (which are more likely to overestimate *L*_∞_) whereas otolith data is a standard approach these days, not accounting for the ageing method in temporal comparisons might give an impression of increasing *K* and declining *L*_∞_ through time, when in fact this might be simply due to different methods used. Given that fisheries and global heating is predicted to increase early growth and reduce asymptotic lengths in many fish species (e.g. Audzijonyte et al., 2016), it is especially important that any trends detected are statistically rigorous and account for potential biases due to methodology. When ageing source in long term studies is known and there is sufficient overlap in the methods through time, differences in methods could be explicitly accounted for in statistical analyses. This will only be possible when at least some data has been aged using both methods, i.e. statistical correction will not work when all old data is from scales and all new data is from otoliths, as there is insufficient information to assess discrepancies between methods. In such cases specific studies might have to be conducted where fish are aged using both methods (as in Pilipaityte et al., submitted after revisions), correction might potentially be introduced using information from other populations or closely related species, or conclusions about temporal or spatial variation should be made with caution and account for potential over- and under-estimation.

## Conclusion

In this study, we demonstrated that the ageing method significantly influences growth parameter estimates, with scale-derived ages tending to overestimate *L*_∞_ and underestimate *K* compared to otolith-derived estimates. These findings are consistent with previous research and underscore the importance of explicitly accounting for ageing method in comparative analyses. We strongly recommend that all fish growth data provided to FishBase, and other databases include ageing source information as an essential part of results. Future research should also explore species-specific ageing biases and potential correction factors that could be used to harmonise historical and contemporary datasets. By acknowledging and addressing these methodological biases, we can enhance the accuracy of growth parameter estimates, improve their comparability, and support more effective fisheries assessment and management.

## Acknowledgements

We would like to thank Fish Ecology Lab staff (Audrius Steponėnas, Linas Ložys, Justas Dainys) and everyone involved in fish data collection and digitisation. This study was supported by the State Scientific Research Institute Nature Research Centre PhD scholarship.

## Author contributions

This study was conceived and designed by A.A, E.P and E.J. Curonian Lagoon fish species data were collected by Ž.P. and E.P. Fish age estimation was done by E.P. and E.J. Statistical analysis was conducted by E.P., in consultation with A.A. and E.J. All authors contribute to the manuscript preparation.

## Data availability

All data and R code necessary to reproduce the results presented in this manuscript will be made publicly available upon article acceptance. Data and R code are available on GitHub:

## Ethical approval

Not applicable.

